# eRNAs Modulate mRNA Stability and Translation Efficiency to Bridge Transcriptional and Post-transcriptional Gene Regulation

**DOI:** 10.1101/2025.07.06.663389

**Authors:** Rene Kuklinkova, Natalia Benova, Chinedu A. Anene

## Abstract

Enhancer RNAs (eRNAs) are best known for their role in transcriptional regulation, where they facilitate enhancer-promoter communication and chromatin remodelling. Yet growing evidence suggests that their function may extend beyond the nucleus. Here, we systematically characterise the decay kinetics of eRNAs across human cell types using time-resolved transcriptomics and kinetic modelling. While most eRNAs undergo canonical exponential decay, a subset displays non-linear dynamics, suggesting context-dependent degradation mechanisms. Perturbation of core decay regulators, including components of the m^6^A and CCR4-NOT pathways, reveals that eRNA stability is modulated by a patchwork of pathways governing mRNA turnover. Integrating transcriptome-wide ribosome profiling, RNA-Seq, and half-life data, we identify eRNAs associated with changes in mRNA stability and translation efficiency of their target protein-coding transcripts. Functional validation of one such eRNA, en4528, shows it regulates CDKN2C mRNA independently of transcription and impacts cell migration. These findings redefine the regulatory scope of eRNAs, positioning them as active participants in post-transcriptional gene control and cellular behaviour. The resulting decay profiles and regulatory annotations have been incorporated into the eRNAkit database, available at https://github.com/AneneLab/eRNAkit, enhancing its capacity for integrative systems-level analysis of eRNA function.

## Introduction

eRNAs are a heterogenous class of non-coding RNAs transcribed from active enhancer regions and have traditionally been studied in the context of transcriptional regulation. Initially thought to be transcriptional byproducts, eRNAs are now recognised as active molecules that facilitate enhancer-promoter communication, promote chromatin remodelling, recruit transcriptional co-activators, and sequester repressive factors ^1–3^. These genomic cis-acting roles have positioned eRNAs as central players in enhancer-mediated control of gene expression, particularly during development, differentiation, and stress responses ^4,5^. While most studies have focused on nuclear functions of eRNAs, a growing body of evidence points to their widespread subcellular localisations and potential post-transcriptional functions ^6–8^. Notably, some eRNAs have been detected in the cytoplasm compartments (e.g., mitochondria and membrane), where they interact with mature mRNAs ^6^. These interactions raise the possibility that eRNAs may act in trans to modulate the fate of target mRNAs, challenging the conventional view of enhancer activity as exclusively transcriptional.

Post-transcriptional regulation mechanisms, particularly control over mRNA stability and translation efficiency shape global gene expression. mRNA half-life determines transcript persistence, while translation efficiency (TE) governs the rate of protein synthesis per mRNA molecule ^9,10^. Both are highly dynamic and responsive to physiological cues. They are typically regulated by RNA sequence features, secondary structures, RNA binding proteins, and microRNAs ^11,12^. However, whether eRNAs contribute to these layers of regulation remains largely unexplored. Our recent large-scale analysis of eRNA localisation and interactions hint that subsets of eRNAs may influence post-transcriptional processes ^6^: some co-sediment with polysomes, sponge miRNAs or interact with mRNAs in a translation dependent manner. Moreover, RBP-dependent mechanisms modulate eRNA-mRNA interactions. These findings raise the possibility that eRNAs may modulate mRNA stability or translation by forming direct RNA–RNA duplexes or acting as scaffolds for regulatory complexes. Yet, comprehensive analyses that link eRNA–mRNA interactions to measurable effects on RNA decay or translation are lacking.

To address this gap, we combine transcriptome-wide maps of eRNA-mRNA interactions with RNA decay kinetics and ribosome profiling data across human cell types. We test the hypothesis that eRNAs modulate mRNA stability and translation efficiency, either by stabilising transcripts, enhancing translation, or both. By integrating localisation, expression, and transcription inhibition assays, we identify eRNAs that predict changes in target mRNA half-lives and ribosome occupancy, suggesting functional relevance for these interactions. Targeted in-vitro knockdown of a candidate eRNA validates these findings, revealing transcription-independent effects on mRNA stability and translation efficiency.

Our findings suggest that enhancers do more than initiate transcription or shape chromatin, they can also influence gene expression after transcription has occurred. We propose a model in which eRNAs play a post-transcriptional role by affecting both mRNA stability and translation. This dual function highlights a more dynamic, RNA-driven view of enhancer activity, positioning eRNAs as important regulators that bridge transcriptional and post-transcriptional gene regulation.

## Results

### Kinetic modelling reveals eRNA stability landscape

Transcription at enhancer regions is a hallmark of active enhancers ^13,14^, and the involvement of eRNAs in cis-regulatory gene expression is well documented ^2^. However, despite evidence that eRNAs localise to all major subcellular compartments and interact with other RNAs ^6,7^, their genome-wide decay dynamics remain largely unexplored. Characterising eRNA stability is the first step to linking them to post-transcriptional regulation of target mRNAs. To address this, we compiled a large dataset of RNA-Seq decay experiments based on time-course Actinomycin D treatment, spanning diverse cell types and conditions (Table S1). We leveraged our previously curated consensus annotation of eRNAs ^6^, generated across multiple cell types, to enable systematic analysis of eRNA stability across these datasets.

To estimate eRNA decay rates, we applied both first-order and second-order kinetic models to the time-course datasets (Methods). The first-order model assumes a constant exponential decay, while the second-order model allows for more complex, nonlinear decay behaviour. Combining dual-model approach enabled fine-mapping of decay kinetics across eRNAs. We fitted both models across 92 complete datasets (Table S2). For each eRNA in each dataset, we evaluated model fit using ANOVA (p < 0.05) and the Akaike Information Criterion (AIC) to determine which model better captured the underlying decay dynamics.

Across most datasets, the majority of eRNAs that passed quality filtering were best described by a first-order exponential decay model (mean: 95%; range: 81–99%), suggesting that classical decay kinetics predominate at the transcriptome level. Despite this global trend, eRNA half-lives exhibited substantial variability both across transcripts and between cell types, while remaining highly consistent between biological replicates within each cell line (Figure 1a, Table S3). To investigate the molecular pathways governing eRNA decay, we compared half-life distributions between control conditions and perturbations of known RNA decay regulators. Surprisingly, knockdown of YTHDF2 or METTL3, key components of the m^6^A-dependent decay pathway, did not lead to global changes in eRNA stability in either C4-2 or HeLa cells (Figure 1b-c). In HepG2 cells, knockdown of IGF2BP1 and IGF2BP2, RNA-binding proteins known to stabilise mRNAs by counteracting decay, resulted in a consistent increase in global eRNA half-lives (Figure 1d), suggesting a previously unappreciated role for these factors in eRNA turnover. IGF2BP3 depletion, by contrast, had no detectable effect (Figure 1d). Meanwhile, knockdown of RNF219, a ubiquitin ligase that modulates CCR4-NOT-mediated deadenylation, also increased eRNA decay in HeLa cells (Figure 1e), whereas knockdown of CNOT3, a core component of the CCR4-NOT complex, paradoxically enhanced eRNA degradation in CCRF-CEM cells (Figure 1f). These context-dependent outcomes highlight that some eRNA turnover may be governed by distinct, possibly non-canonical decay pathways, differing from those established for mRNA biology.

**Figure 1:**
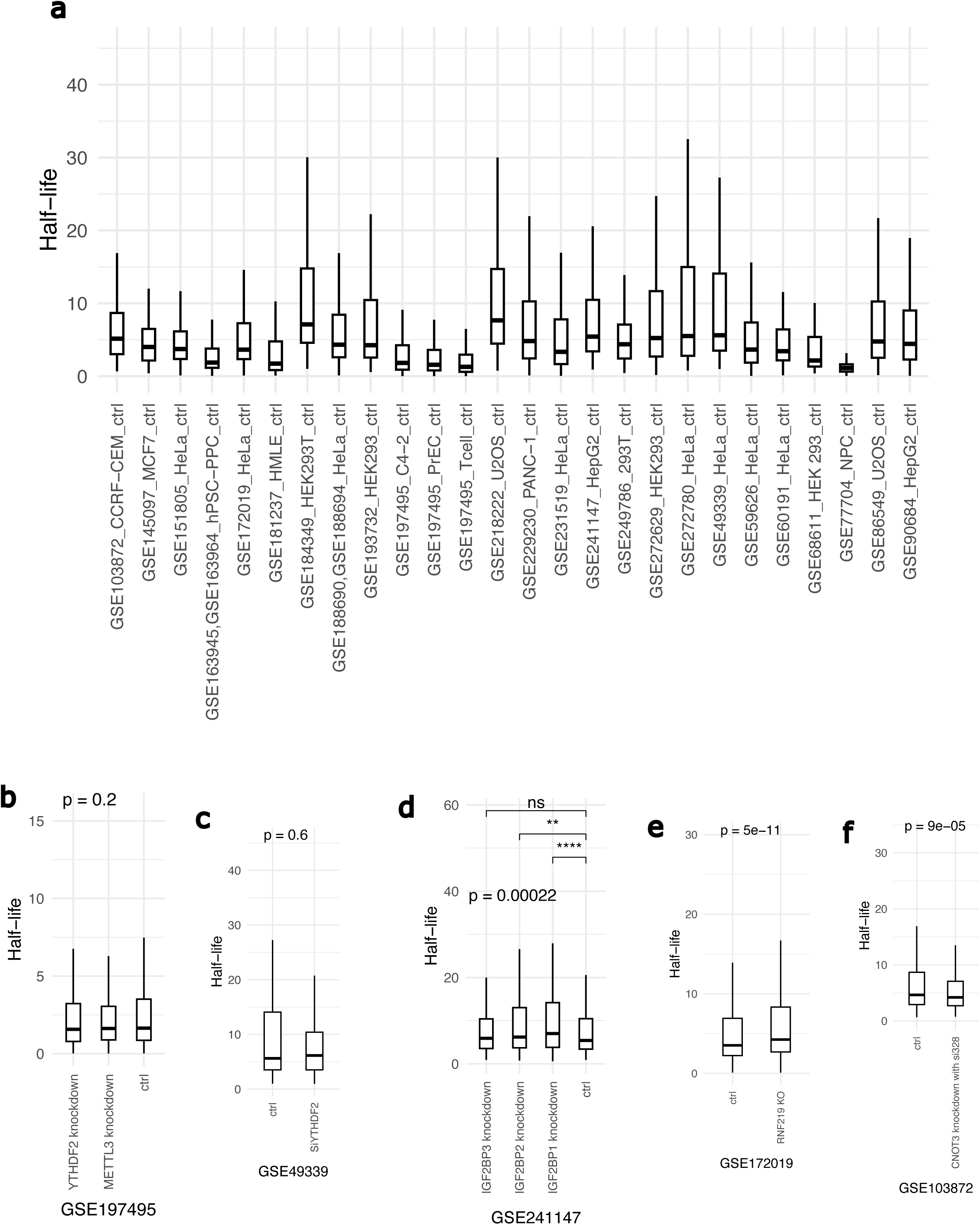
eRNA decay kinetics under basal and perturbed conditions across cell types. (a) Distribution of eRNA half-lives across all high-quality control samples from the compiled Actinomycin D time-course datasets. Each point represents the estimated half-life of an individual eRNA, measured under basal (untreated) conditions. (b-f) Comparative distribution of eRNA half-lives between control and knockdown (or knockout) conditions for key RNA decay regulators across distinct cell lines. (b) Knockdown of YTHDF2 and METTL3, core components of the m^6^A-dependent decay pathway (cell line: C4-2). (c) Isolated effect of YTHDF2 knockdown versus control (cell line: HeLa). (d) Knockdown of IGF2BP1, IGF2BP2, and IGF2BP3 compared to control, testing the impact of this mRNA-stabilising protein family on eRNA stability (cell line: HepG2). (e) Knockout of RNF219, a CCR4–NOT–associated E3 ubiquitin ligase, compared to control (cell line: HeLa). (f) Knockdown of CNOT3, a scaffold protein of the CCR4-NOT complex, compared to control (cell line: CCRF-CEM). P-values were computed using either Wilcoxon rank-sum test or Kruskal-Walli’s test, as appropriate. Boxplots show median (centre line), interquartile range (box), and 1.5× IQR whiskers; outliers are not shown.

### eRNAs mirror mRNA decay kinetics, with minor deviations

To compare eRNA decay patterns with genes, we applied the same model selection approach to genes. As expected from prior studies, the simpler first-order exponential decay model provided the best fit for most mRNAs, consistent with canonical decay kinetics (Table S4). When comparing model selection outcomes between eRNAs and gene transcripts, we observed a small but statistically significant difference: the second-order model was selected slightly more frequently for genes than for eRNAs (mean: 5.5% vs. 4.94%; *p* = 0.011; Figure 2a). A similar trend was observed when restricting the analysis to protein-coding genes (5.77%; *p* = 0.013). While these differences are modest in magnitude, they may reflect subtle distinctions in decay dynamics between eRNAs and traditional mRNAs. As most transcripts (95%) were well-fitted to a first-order decay model (see Methods), we used half-life estimates derived from this model for all subsequent analyses.

**Figure 2:**
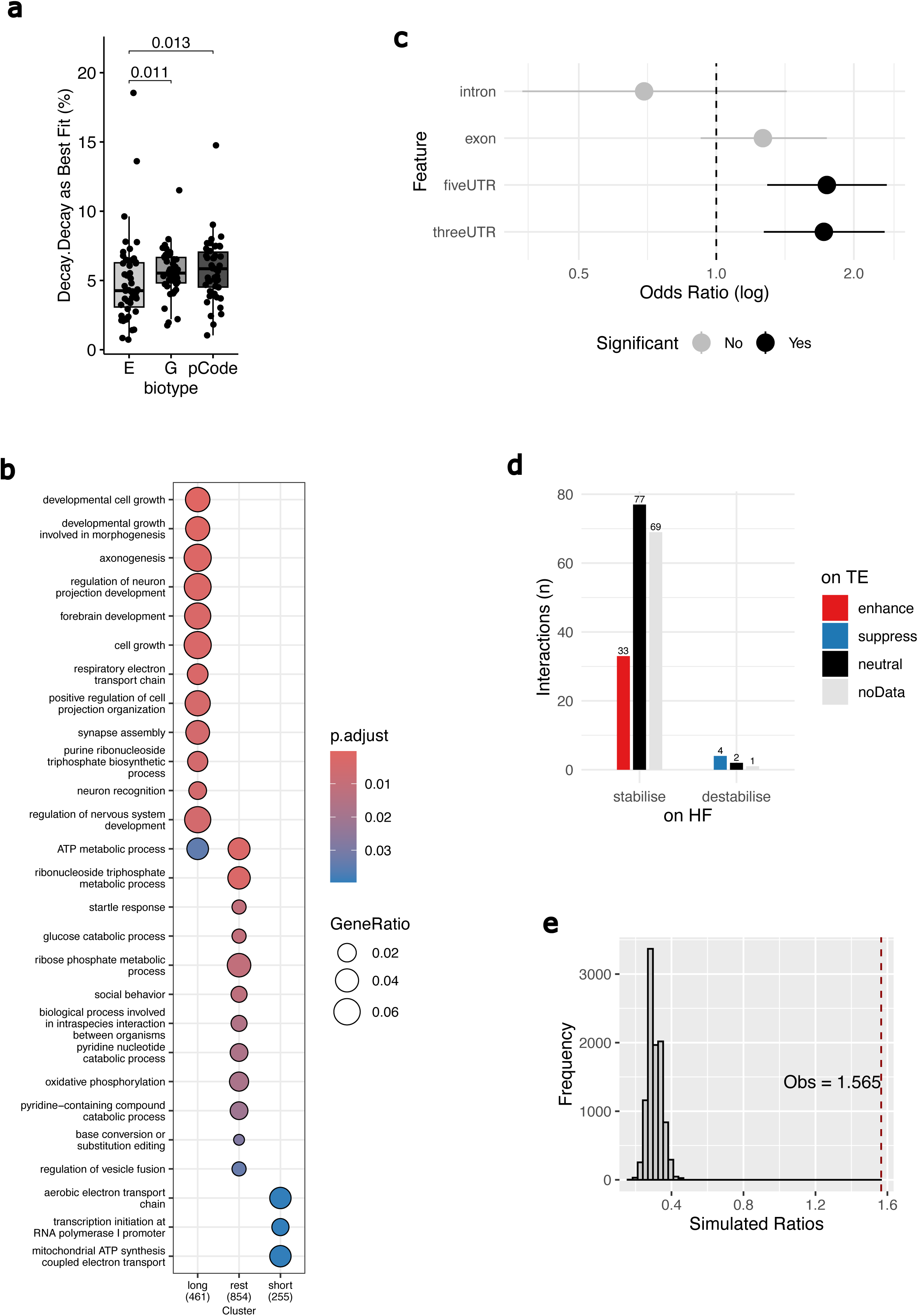
Features and functional associations of eRNA decay kinetics and regulatory potential. (a) Boxplot comparing the percentage of transcripts whose decay profiles are best explained by a second-order (nonlinear) decay model across different RNA types. E: eRNAs, G: all genes, pCode: protein-coding genes. P-values were calculated using the Wilcoxon rank-sum test. (b) Gene Ontology enrichment analysis of mRNAs targeted by eRNAs from distinct half-life classes (short, rest, and long). Top 12 enrichment is shown for biological process terms with reduced redundancy via semantic similarity filtering. (c) Forest plot showing odds ratios and 95% confidence intervals for enrichment of sequence features (5′UTR, 3′UTR, exon, or intron) at the interaction sites of regulatory eRNA–mRNA pairs versus non-regulatory pairs. Annotations are based on canonical transcript features overlapping the mRNA-side of each interaction. (d) Number of eRNAs identified as regulatory based on changes in target mRNA half-life, translation efficiency (TE), or both. (e) Distribution of overlap ratios derived from expression-shuffling simulations comparing half-life- and TE-associated regulatory eRNAs. The vertical line marks the observed number of observed overlapping regulatory eRNAs.

### eRNA kinetics stratifies target mRNAs into distinct regulatory programs

Genes with related functions are often co-regulated such that their mRNAs have similar half-lives ^9,10^. To investigate whether eRNA half-lives play a comparable post-transcriptional regulatory role, we grouped eRNAs into stability classes (short-, rest-, and long-lived) based on half-life thresholds (Methods). We then leveraged experimentally derived RNA-RNA interaction datasets (KARR-Seq, RIC-Seq and PARIS), spanning diverse cell types and conditions ^6^ to link putative target mRNAs for each eRNA. Across all datasets, we identified 1043 eRNAs with consistent stability classification (i.e., assigned to the same group in ≥60% of the datasets), representing 6780 eRNA-mRNA interaction pairs (short = 1059; long = 1263, rest=4458). Gene ontology enrichment analysis of each group revealed distinct patterns of functional associations (Figure 2b, Table S5). Targets of the long-lived eRNAs were enriched for developmental and morphogenetic processes, including those linked to cell growth and tissue organisation. This is consistent with the notion that such processes require prolonged and coordinated gene regulation, suggesting that eRNA stability may be a key factor in supporting long-term post-transcriptional control. In contrast, short-lived eRNAs, though fewer GO terms, showed focused enrichment in mitochondrial electron transport and transcriptional initiation, possibly reflecting dynamic regulatory functions in energy metabolism and cellular adaption. The rest group, comprising eRNAs with half-lives in the interquartile range (25^th^-75^th^ percentile), displayed a broader and more functionally diverse enrichment profile. In addition to ATP and nucleotide metabolism, oxidative phosphorylation, and glucose catabolism, these targets were also enriched for processes involved in vesicle trafficking, intercellular communication, and behavioural responses. Restricting the analysis to eRNA-mRNA interaction pairs without known genomic associations (cis or enhancer-promoter) did not alter the enrichment patterns (Figure S2a, Table S6), indicating that these functional associations are unlikely to result from shared chromatin proximity alone. Together, these findings support the hypothesis that eRNAs exert post-transcriptional regulatory functions, with half-life–linked RNA-RNA interaction patterns reflecting distinct modes of regulation across biological pathways.

### Stability of target mRNAs reflects regulatory influence of eRNAs

Having shown that eRNAs with different stability profiles are associated with distinct functional pathways through their target mRNAs, and that these associations are independent of known genomic interactions. We next asked whether specific eRNAs contribute to the regulation of mRNA turnover, consistent with the hypothesis that their decay properties may impact those of their targets. To this end, we stratified samples (each representing a complete time course) into short and long-lived groups for each eRNA, using the median half-life across all samples. Using the same RNA-RNA interaction pairs as above, we then compared the half-lives of target mRNAs between these eRNA-defined groups using the Wilcoxon rank-sum test.

Of 14,761 eRNA-mRNA interaction pairs tested, 186 pairs showed statistically significant differences in target mRNA half-life at a false discovery rate (FDR) < 0.05 (Table S7). The vast majority (179) exhibited increased mRNA stability associated with stable eRNA groups, while only 7 pairs showed decreased stability. None of the destabilising eRNAs had multiple significant target pairs, whereas among the unique stabilising eRNAs 18 out of 155 with significant target interactions had multiple significant pairs, and in all such cases, the direction of effect was consistent across targets. The top three eRNAs with the highest number of significant interactions had 6, 6, and 5 significant pairs respectively (Figure S3a-b, Table S7), a pattern not solely explained by the total number of tested pairs per eRNA, indicating potential key regulatory roles for these eRNAs. While 14,761 unique eRNA-mRNA pairs were tested, these involved 2,614 distinct eRNAs. Of these, 155 eRNAs (approximately 6%) had at least one target pair showing a significant effect on mRNA half-life, indicating that a substantial subset of eRNAs may play a regulatory role in mRNA stability. Supporting this notion, we examined their subcellular localisation in K562 and HepG2 cells using the eRNAkit resource ^6^ and found that all 155 unique eRNAs were detected in cytoplasmic fractions, including insoluble and membrane compartments, across polyA+, polyA−, and total RNA samples (Figure S1a). These findings support a model in which a distinct subset of eRNAs engages in post-transcriptional regulation, with their cytoplasmic localisation aligning with a potential role in modulating mRNA stability.

### Regulatory eRNA-mRNA interactions are enriched in untranslated regions

RNA structure can influence RNA-RNA interactions and affect transcript stability ^15,16^. To determine whether specific mRNA features are associated with the observed regulatory effects of eRNAs on mRNA stability, we examined the overlap of annotated mRNA regions with the precise sites of the eRNA-mRNA interactions. We compared regulatory pairs (those with significant effects on target mRNA stability) against non-regulatory pairs using Fisher’s exact tests. We observed a significant enrichment for interactions overlapping 3′ untranslated regions (3′UTRs; odds ratio (OR) = 1.85, 95% confidence interval (CI) = 1.29–2.66, p = 4.8 × 10⁻⁴) and 5′ untranslated regions (5′UTRs; OR = 1.91, 95% CI = 1.33–2.74, p = 2.9 × 10⁻⁴) among regulatory pairs (Figure 2c). Overlap with exonic regions showed a non-significant trend (OR = 1.42, p = 0.061), while intronic regions showed no significant association (OR = 1.47, p = 0.33). These findings indicate that eRNA-mediated modulation of mRNA stability preferentially occurs at untranslated regions, consistent with the functional importance of these regions in post-transcriptional regulation.

### Regulatory eRNA-mRNA interactions coordinate mRNA stability and translation efficiency

The enrichment of UTRs among regulatory eRNA-mRNA interactions prompted us to investigate whether these interactions also influence translation efficiency (TE). Given the well-established role of UTRs in modulating translation, we reasoned that eRNAs affecting mRNA stability might also exert coordinated effects on translation. To investigate this, we examined TE changes for mRNAs involved in regulatory eRNA-mRNA pairs using large-scale ribosome profiling datasets from RPFdb ^17^. For each eRNA, we stratified samples into high- and low-expression groups based on the median expression level across all samples. We then compared target mRNA TE between these groups to assess whether eRNA levels are associated with consistent changes in translation, in line with their observed effects on mRNA stability. Strikingly, 37 of 186 (19.9%) regulatory eRNA-mRNA pairs showed significant TE changes (FDR < 0.05), and in all cases, the direction of the TE change matched that of the stability effect: eRNAs associated with increased mRNA stability also enhanced TE, while those linked to destabilisation suppressed TE (Figure 2d, Table S7).

To assess whether this pattern could arise by chance, we performed two sets of simulations: (1) shuffling eRNA expression while preserving target pairings and TE data, and (2) permuting eRNA-mRNA interaction pairs while retaining expression and TE data. The observed number of concordant TE changes significantly exceeded expectations under both null models: the empirical p-value for the expression shuffle was 0 (Figure 2e), and for the name shuffle was 0.0575 (Figure S4a). These results reinforce that the association between eRNA expression and target mRNA TE is not a product of random pairing or expression variation alone. Together, these observations support a model in which eRNAs not only stabilise or destabilise their target transcripts, but also influence their translational output, potentially amplifying their regulatory impact at the protein level.

### A Cytoplasm-Localised eRNA Regulates Target Stability and Cellular Migration

To experimentally validate our model of eRNA-mediated post-transcriptional regulation, we selected a representative eRNA-mRNA interaction pair, en4528 and CDKN2C, for functional interrogation. This eRNA exhibited cytoplasmic localisation and formed RNA-RNA contacts with three protein coding mRNAs ^6^. Of these, CDKN2C showed a statistically significant increase in mRNA stability associated with the stable en4528 group, while the other two mRNAs, DHDDS and C1orf185, did not show a significant effect in the stability analysis.

To specifically assess the functional contribution of the cytoplasmic pool of the eRNA, we employed siRNA-mediated knockdown directed through the RNA-induced silencing complex (RISC), thereby selectively depleting cytoplasmic en4528 levels ^18^. qPCR confirmed efficient depletion of the eRNA (Figure 3a). We next evaluated the expression of the regulated mRNA target following siRNA knockdown. Consistent with the stability analysis, CDKN2C was significantly downregulated upon cytoplasmic depletion of en4528 (Figure 3a), supporting a post-transcriptional stabilising role for the eRNA. This regulatory effect was not reversed by pre-treatment with actinomycin D (Figure 3b), confirming that it occurs independently of transcriptional input. Similarly, FAF1, a gene located within the same topologically associating domain (TAD) and locus ^18^, also exhibited significant downregulation under the same conditions (Figure 3a). Like CDKN2C, this effect was not resolved by transcriptional blockade (Figure 3b), indicating that FAF is also regulated post-transcriptionally by en4528. These findings indicate that additional regulatory eRNA-mRNA pairs may exist, which will require further investigation to resolve.

**Figure 3:**
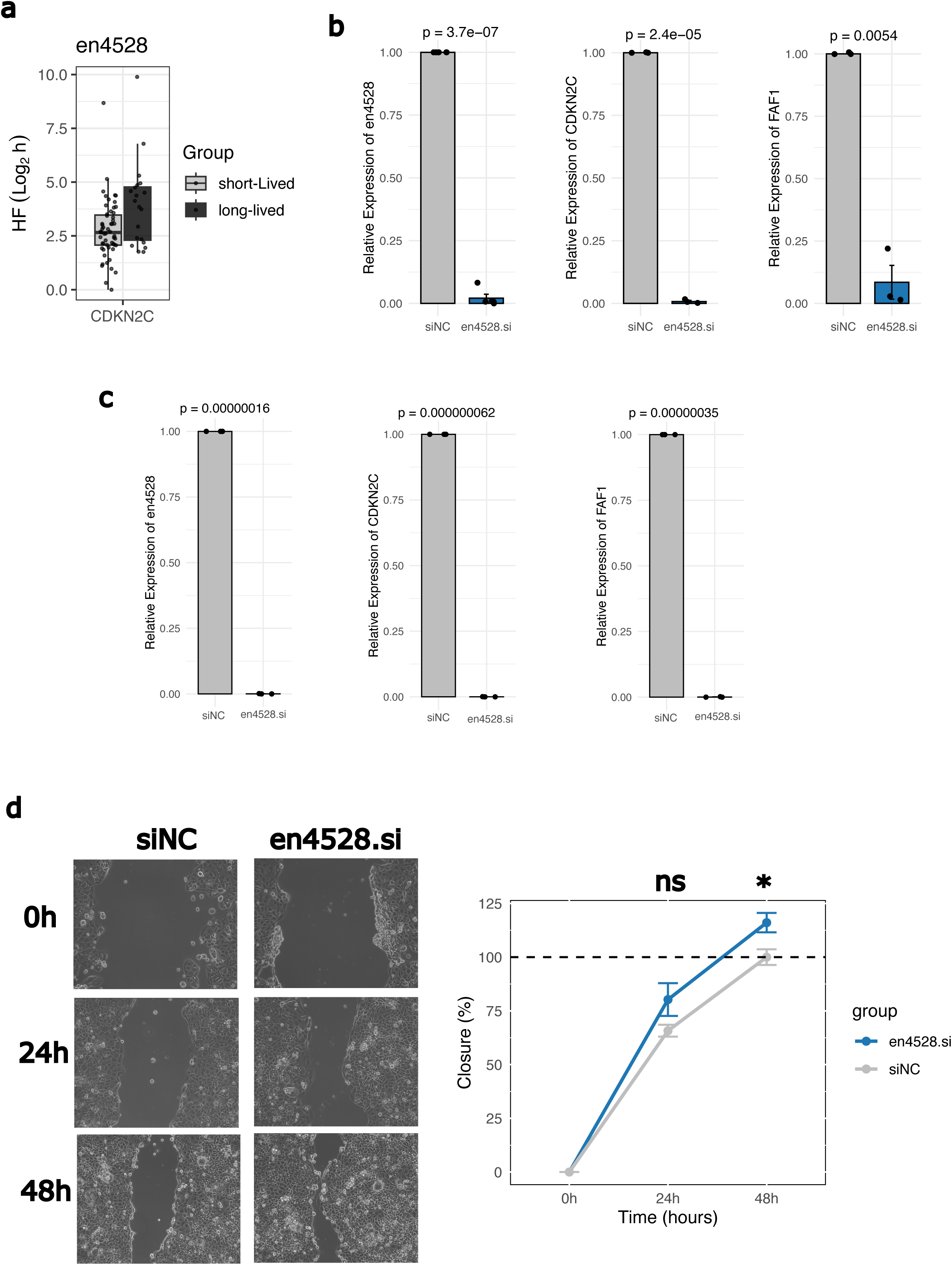
Functional validation of eRNA en4528 and its regulatory impact on target mRNAs and cell migration. (a) Comparison of CDKN2C mRNA half-life between groups of samples classified by short-lived versus long-lived en4528 in the main time-course dataset. (b-c) Relative expression levels of en4528, CDKN2C, and FAF1 measured by RT-qPCR following (b) siRNA-mediated knockdown of en4528 (en4528.si) compared to non-targeting control (siNC) at 48 hours (c) pretreatment with actinomycin D for 30 minutes before siRNA-mediated knockdown of en4528 (en4528.si) compared to non-targeting control (siNC) at 2 hours. GAPDH was used as a loading control. Data are shown as mean ± SEM from at least three independent experiments. (d) Scratch wound migration assay assessing the effect of en4528 knockdown on cell motility. Left panel shows representative images at 0-48-hours post-scratch for en4528.si and non-targeting control cells. Right panel quantifies percentage wound closure over time (mean ± SEM from at least three independent experiments).

Finally, to examine the phenotypic consequence of this regulatory axis, we performed a wound healing (scratch) assay following siRNA depletion of en4528. We observed a significant increase in cell migration compared to control conditions (Figure 3c), indicating that loss of the eRNA alters cellular function in a manner consistent with its inferred post-transcriptional regulatory role. Together, these findings provide experimental support for the functional relevance of post-transcriptional eRNA-mediated regulation of mRNA stability, highlight the complexity of eRNA-target interactions within shared chromatin environments, and demonstrate that such interactions can elicit measurable phenotypic outcomes.

## Discussion

Despite growing evidence that eRNAs are essential for enhancer function, establishing direct mechanistic links between eRNAs and specific regulatory outcomes remains a key challenge in RNA biology ^3,19^. While previous work has focused predominantly on transcriptional effects at nearby promoters, whether eRNAs exert broader post-transcriptional influence, particularly through control of mRNA stability, has remained largely unexplored. Here, we present a systematic framework for characterising the decay kinetics of eRNAs and their potential regulatory impact on target transcript stability and translation efficiency. Using time-resolved RNA decay datasets and kinetic modelling, we provide the first transcriptome-wide map of eRNA half-lives across diverse cell types. We show that most eRNAs follow canonical exponential decay, consistent with classical RNA turnover ^20,21^. However, we also identified a subset of eRNAs with atypical decay profiles. These deviations may reflect the influence of additional layers of regulation, such as dynamic RNA-binding protein (RBP) interactions, alternative degradation pathways, or context-dependent RNA modifications. To explore the underlying mechanisms, we evaluated half-lives under perturbations of key post-transcriptional regulators. Intriguingly, silencing canonical decay factors such as YTHDF2 and METTL3, central to the m^6^A-mediated RNA decay pathway ^22^, did not result in global changes in eRNA stability. This implies that m^6^A modification, while central to mRNA decay, may play a limited role in regulating eRNA stability.

Knockdown of IGF2BP family members, known to stabilise mRNAs by antagonising decay pathways ^23^, unexpectedly led to increased eRNA half-lives. This counterintuitive outcome, together with the contrasting effects observed upon perturbation of RNF219 and CNOT3, both associated with the CCR4-NOT deadenylation complex ^24^, suggests that eRNA stability is governed by a more intricate and possibly distinct regulatory logic than that of canonical mRNAs. Building on this, we propose a model in which eRNA turnover is governed by a patchwork of RBP-dependent regulatory mechanisms, some of which may be cell type- or context-dependent. Moreover, the involvement of cytoplasmic decay factors also supports the functional relevance of eRNAs beyond the nucleus and argue for a broader reframing of eRNA function and fate. While many eRNAs may be rapidly turned over in a constitutive manner, others appear to be selectively regulated, potentially enabling them to exert fine-tuned control over gene expression. The existence of such regulated decay could serve as a mechanism to temporally constrain or amplify the impact of eRNAs under cellular states or signalling conditions. As such, understanding the determinants of eRNA stability may prove key to elucidating their full spectrum of regulatory functions.

While eRNAs are well-recognised for their activity at enhancers and in the nucleus, their presence in the cytoplasm and their capacity to form RNA-RNA interactions raise the possibility of a broader post-transcriptional role. Through comparative analysis of target mRNA decay rates across eRNA-defined stability groups, we identify a subset of eRNAs (approximately 6% of those tested), that exert significant, directional effects on mRNA stability.

An intriguing observation from this analysis is that 96% of eRNAs with significant post-transcriptional associations were linked to increased mRNA stability, and this pattern was mirrored at the level of translation efficiency. eRNA that stabilise their targets also increased ribosome occupancy on those targets. This consistent asymmetry suggests a directional bias in eRNA-mediated post-transcriptional regulation. Rather than acting as a double-edged sword, some eRNAs appear to reinforce the gene expression programmes they help initiate. This is conceptually consistent with the fundamental role of enhancers as activators, “enhancers finish the job they start.” That is, having promoted transcriptional activation, a subset of eRNAs may sustain expression post-transcriptionally by stabilising their target mRNAs. Destabilising the very transcripts, they help produce would represent a paradoxical mode of regulation. Instead, our data suggest that some eRNAs act to consolidate and prolong gene expression outputs, adding a layer of temporal reinforcement to the enhancer’s initial activation signal.

Given the conservative nature of our curation and filtering steps in eRNAkit ^6^, which prioritised high-confidence eRNA-mRNA interaction pairs, this proportion likely underestimates the true extent of the consolidate regulation. Supporting this view, exploratory analysis using a less stringently filtered dataset suggests the existence of additional putative regulatory interactions (data not shown). The observed enrichment of UTRs among these regulatory eRNA-mRNA interactions provided further support for regulatory functions beyond the nucleus. UTRs are known hubs of post-transcriptional control, harbouring binding sites for RNA-binding proteins and miRNAs, as well as structural elements affecting RNA stability and translation ^25^. eRNAs could exploit these domains to modulate mRNA decay or engage canonical RNA decay machinery. Prompted by this, we examined whether eRNAs that modulate stability also influence the translational output of their targets. Strikingly, nearly 20% of the stability-associated eRNA-mRNA pairs exhibited significant changes in translation efficiency (TE), and in all cases, the direction of TE change mirrored the stability effect. This concordance suggests that eRNA-mediated stabilisation of target mRNAs not only preserves transcript abundance but may also enhance their engagement with the translational machinery, amplifying the regulatory impact at the protein level.

To evaluate whether the coupling between eRNA-dependent target stability and translation efficiency could arise by chance, we implemented two null models based on expression and interaction shuffling. While the empirical p-value from the eRNA expression shuffle was 0, indicating a highly non-random association, the name-shuffling yielded a borderline value (p = 0.0575). We interpret this result cautiously: because the null model was generated by permuting known eRNA–mRNA pairs, it may have inadvertently retained genuine but unannotated interactions. Given the complexity and redundancy of RNA-RNA regulatory networks, such spurious retention could inflate the null distribution and reduce statistical resolution. Accordingly, the name-shuffling simulation likely represents a conservative test of enrichment, and the consistency of our findings across both models supports the biological relevance of coordinated regulation between stability and translation.

Collectively, our results propose a model in which a subset of cytoplasm-localised eRNAs function as trans-acting regulators of mRNA turnover and translation, thereby extending the functional repertoire of enhancers beyond the nucleus. These eRNAs may act through direct base-pairing interactions or via recruitment of decay- or translation-associated complexes. Importantly, our final experimental validation demonstrates that perturbation of a cytoplasm localised eRNA affects both target stability and cell migration, supporting the idea that such post-transcriptional regulation can drive phenotypic outcomes.

While our study establishes a framework for mapping eRNA decay kinetics and linking them to target mRNA regulation, several limitations remain. First, although we leveraged a large and diverse set of time-resolved RNA-Seq datasets, Actinomycin D-based inhibition, may not distinguish between nuclear retention and true decay. Second, our identification of regulatory eRNA-mRNA pairs relied on conservative high-confidence annotations. While this conservative approach minimizes false positives, it may underestimate the full extent of eRNA-mediated regulation, especially for condition-specific or transient interactions. Third, although we show that eRNA-mediated regulation preferentially targets UTR regions and that changes in stability frequently coincide with changes in translation efficiency, the mechanistic basis for these effects remains to be fully elucidated. Whether eRNAs act through direct base-pairing, recruitment of specific RNA-binding proteins (RBPs), or structural remodelling of target mRNAs remains an open question. Finally, further functional studies, including direct binding assays and reporter-based validation across multiple eRNA-mRNA pairs, are needed to confirm the generality of our findings and uncover causal mechanisms. The extent to which these regulatory interactions contribute to physiological outcomes, particularly in disease contexts such as cancer or neurodegeneration, also warrants future exploration. These limitations notwithstanding, this study provides a conceptual and experimental framework for decoding eRNA function beyond cis-regulatory transcription.

Future work will be required to resolve the mechanisms underlying eRNA selectivity, to dissect their interaction partners, and to explore the broader physiological relevance of eRNA-mediated post-transcriptional networks in development and disease. Notably, the core findings from this study have been incorporated into the eRNAkit resource, expanding its utility for integrative systems biology analyses of eRNA function across regulatory layers.

## Methods

### eRNA Annotation and RNA-RNA Interaction Curation

The eRNA annotation across multiple cell types and the compilation of high-confidence RNA-RNA interactions (from KARR-Seq, RIC-Seq, and PARIS datasets) used in this study were previously described in ^6^.

### RNA Stability Modelling Compilation of RNA decay datasets

To characterise RNA stability transcriptome-wide, we manually curated RNA-Seq datasets from actinomycin D (ActD) time-course experiments, which inhibit transcription and allow estimation of RNA decay rates. Public datasets were obtained from GEO and SRA and screened for appropriate time-point coverage, metadata clarity, and experimental consistency. We compiled 364 samples across 16 human cell lines, forming 96 complete time-course datasets, each comprising at least three post-treatment time points ^23,26–48^. Of these, 27 datasets represent wild-type (unperturbed) cells, while the remaining 69 capture a wide range of genetic perturbations, with a particular focus on knockdown of RNA decay-associated proteins. These included core components and modulators of known decay pathways, enabling systematic evaluation of stability regulation under both basal and perturbed conditions. These compiled 96 complete datasets are herein referred to as the time-course datasets and form the basis of our transcriptome-wide RNA stability analysis. Detailed sample-level information is provided in Supplementary Table S1 and S2.

Raw RNA-Seq reads corresponding to these datasets were obtained from SRA and processed using our established pipeline ^49^. Reads were quality filtered (Q < 20) and adapter trimmed using Trimmomatic (v0.39) ^50^, then aligned to the human reference genome (hg38) using HISAT2 (v2.1.0) with default settings ^51^. Gene level and eRNA counts were quantified using HTSeq (v0.11.1)^52^ against our previously curated consensus eRNA annotation or human GRCH38 annotation (Encode v111 release). The expression levels were normalised by CPM.

### Kinetic modelling of RNA decay

To characterise the decay kinetics of RNAs across the datasets, we applied both first-order and second-order exponential decay models to the expression profiles of each transcript per time-course dataset. Only transcripts with detectable expression (CPM > 1 in ≥2 time points) across at least one complete dataset were retained. For each eRNA and each time course, we modelled the decay of expression levels over time using the following formulations:

Standard exponential decay (first-order form):

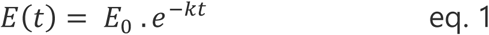

where:

*t*: *Time*

*E*_0_: *Initial expression at t* = 0

*E*(*t*): *Expression at t* 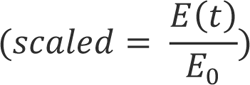

*e*: *Euler’s number*

*k*: *Decay rate constant*

The half-life was calculated as: 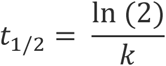

The decay constant *k* was estimated as the slope of the linear regression of expression over time: *k* = *slope of (log (E*(*t*) ∼ *t*)

Decay-of-decay model (second-order form):

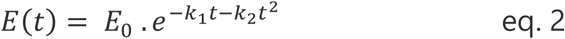

where:
*k*_1_: *Models the primary decay rate;*

*k*_2_: *acounts for curvature in the decay*

The decay constants *k*_1_ and *k*_2_ were estimated by fitting a quadratic regression to the expression values over time, allowing for non-linear decay behaviour not captured by standard exponential fits. For these fits, half-lives were estimated numerically by solving for the time at which *E*(*t*) = 0.5*E*_0_.

For model selection, an ANOVA test was used to evaluate whether the second-order model provided a statistically significant improvement in fit over the simpler first-order model. If the ANOVA result was significant (p < 0.05), the model with the lower Akaike Information Criterion (AIC) was selected. If the ANOVA was not significant, the first-order model was retained by default. This approach ensured that model complexity was only favoured when justified by both statistical significance and improved goodness-of-fit.

### Stratification of eRNAs by stability class to define regulatory programs

To investigate whether eRNA decay kinetics reflect distinct post-transcriptional regulatory programs, analogous to the known coupling between mRNA half-life and functional co-regulation, we stratified eRNAs into three stability classes, short-lived, rest, and long-lived, based on their estimated half-lives from first-order decay models. Only samples derived from unperturbed (wild type) conditions were included to avoid confounding effects from experimental perturbations. Within each time-course dataset, replicate measurements of half-life were first collapsed by computing the median half-life per eRNA per dataset, providing a robust summary for classification. We then stratified the resulting distribution of eRNA half-lives into quartiles: values below the 25th percentile was designated short-lived, those above the 75th percentile as long-lived, and the remaining as rest. These quartile-based thresholds were selected to provide a clear separation between kinetically distinct groups while avoiding overfitting to noise. To assign a global stability class to each eRNA, we determined the proportion of datasets in which it was consistently assigned to a given class. eRNAs were retained if they received the same classification in ≥60% of datasets (with detection in at least three datasets required), ensuring biological reproducibility across cell types.

The final set of consistently classified eRNAs was then used to investigate shared functional properties of their RNA-RNA interaction partners (mRNAs), using gene ontology (GO) enrichment analysis. We employed the ClusterProfiler R package ^53^ along with the human Bioconductor annotation database (org.Hs.eg.db) to identify enriched biological processes at FDR < 0.05, reducing redundancy via semantic similarity analysis ^54^. Since eRNAs lack direct functional annotation, we adopted this guilt-by-association approach, inferring the functional roles of eRNAs from the GO terms enriched among their mRNA targets. This strategy allowed us to test whether eRNAs with similar decay properties tend to engage similar sets of targets and thus may participate in coordinated post-transcriptional regulatory programs.

### Sample-Level Stratification by eRNA Stability for Target mRNA Analysis

To investigate whether differences in eRNA stability predict changes in the stability of their target mRNAs, we stratified time-course samples into eRNA-specific stability groups. For each eRNA, we first computed the median half-life (50^th^ percentile) across all samples in which the eRNA was detected. Samples with estimated eRNA half-life values below this median were classified as short-lived, and those with half-life values equal to or above the median were classified as long-lived. This binary classification allowed us to compare the half-lives of target mRNAs between conditions where the regulatory eRNA was relatively short- or long-lived, respectively. RNA-RNA interaction data were used to define eRNA-mRNA pairs. Only eRNAs with a non-missing median half-life and detectable expression across samples were retained. This per-eRNA stratification was designed to enable robust statistical comparisons of target transcript kinetics in the presence of stable versus unstable eRNA variants, without requiring global consistency across cell types. We compared the half-lives of target mRNAs between the short- and long-lived eRNA groups using the Wilcoxon rank-sum test and computed FDR per eRNA.

### Translation Efficiency Analysis of Regulatory eRNA-mRNA Pairs

To investigate whether eRNA-mediated regulation extends to translation, we assessed translation efficiency (TE) patterns of target mRNAs involved in previously identified regulatory eRNA–mRNA pairs. Pre-computed TE values for human protein-coding genes were downloaded from RPFdb v3.0 ^17^, a curated database of ribosome profiling data spanning multiple cell lines and conditions. To stratify samples by eRNA expression, we first quantified eRNA abundance from matched RNA-Seq data tracks also available in RPFdb. Specifically, we used bigWigAverageOverBed (UCSC tools) to compute average signal over consensus eRNA coordinates, yielding per-eRNA expression estimates across all samples. The mean0 column (average signal excluding zero-coverage bases) was used for downstream classification. For each eRNA, samples were split into high- and low-expression groups based on the median expression value across all samples. Only eRNAs involved in the previously defined regulatory interaction pairs (i.e., those showing a significant effect on mRNA stability) were retained for TE analysis. We compared the TE of target mRNAs between the high- and low-eRNA expression groups using the Wilcoxon rank-sum test and computed FDR per eRNA.

### Permutation Test

To assess whether the number of regulatory eRNA-mRNA pairs exhibiting concordant effects on mRNA stability and translation efficiency (TE) was significantly greater than expected by chance, we performed two non-parametric permutation tests (Expression-shuffle and Pairing-shuffle null models). A pair was defined as concordant if the direction of the TE change between high and low eRNA expression groups matched the direction of the stability effect (e.g., stabilising eRNAs associated with increased TE). The null distributions were created as follows: 1) The observed eRNA expression values were randomly shuffled across samples, while keeping the original eRNA-mRNA interaction pairs and TE measurements intact. For each permutation, samples were reclassified into high- and low-eRNA expression groups based on the shuffled values, and concordant TE effects were recalculated. 2) The eRNA-mRNA pairings were randomly permuted across the set of expressed RNAs, while preserving the eRNA expression and TE matrices. For each permutation, the number of concordant stabilities-TE relationships were recalculated using the permuted pairs. Each permutation model was repeated 10,000 times to generate a null distribution of the number of concordant pairs. The empirical p-value was defined as the proportion of permutations in which the number of concordant TE effects was greater than or equal to the observed count. This one-tailed test estimates the probability of obtaining the observed level of concordance under the null hypothesis of random association.

### Mapping of Interaction Sites to mRNA Features

To determine whether regulatory eRNA-mRNA interactions preferentially localize to specific regions of target transcripts, we annotated the mRNA interaction sites with canonical transcript features using the TxDb.Hsapiens.UCSC.hg38.knownGene database (v3.21).

Genomic coordinates corresponding to the mRNA side of each interaction were intersected with 5′ untranslated regions (5′UTRs), 3′UTRs, exons, and introns, with overlap scored as present (1) or absent (0). The distribution of these annotated features was then compared between regulatory and non-regulatory eRNA-mRNA pairs using Fisher’s exact test.

### Cell culture

Human ovarian carcinoma cells (OVCAR-3) were obtained from the American Type Culture Collection (ATCC) and certified mycoplasma-free. OVCAR-3 cells were maintained in Roswell Park Memorial Institute (RPMI) 1640 medium supplemented with 20% foetal bovine serum (FBS), and 4 mg/ml human insulin. Cells were cultured at 37°C in a humidified incubator with 5% CO_2_. siRNA inhibitors were transfected using Lipofectamine RNAiMAX (Thermo Fisher Scientific, UK), as previously described ^55^. Details of siRNA sequences used in this study are listed in Table S8.

### RNA extraction and RT-qPCR

Total RNA was extracted from en4528.si, or siNC transfected cells using TRIzol reagent (Thermo Fisher, UK) according to the manufacturer’s instructions. cDNA was synthesised from 500 ng of total RNA using the iScript^TM^ cDNA Synthesis Kit (Bio-Rad, UK).

Quantitative PCR was performed on a CFX96 Real-Time PCR Detection System (Bio-Rad, UK). The ΔΔCT analysis method was used, normalised to the GAPDH as a loading control. The Primer sequences are provided in Table S8.

### Cell migration assay

For wound healing assays, a uniform scratch was introduced into confluent monolayers of cultured cells using a 200μl pipette tip, followed by incubation in serum-free medium (SFM). Images were captured at defined intervals over 48 h using an EVOS FL Auto 2 Imaging System (10× objective). Wound closure was quantified using a custom Python script.

### Statistical analysis

Unless otherwise stated, graphical data represent the mean ± standard error of the mean (SEM) from at least three independent experiments. Differences between groups were assessed using Student’s t-test, Wilcoxon rank-sum test, or one-way ANOVA, as specified in the figure legends. Statistical significance was defined as p < 0.05 for single comparisons and as FDR < 0.05 for multiple testing.

## Data availability

All scripts used for modelling eRNA decay kinetics and annotating regulatory eRNA–mRNA interactions are provided as part of the eRNAkit resource, available at https://github.com/AneneLab/eRNAkit. This toolkit, described in a separate work ^6^, includes code for model fitting, statistical testing, and integration of RNA-Seq and ribosome profiling data. Regulatory annotations for eRNA-mRNA interactions generated in this study are also included within the resource.

## Author contributions

Project was conceived of by C.A.A. Data curation and processing were done by C.A.A, N.B, and R.K. C.A.A supervised the project and wrote the manuscript. All authors edited the final manuscript.

## Disclosure and declaration

We declare no conflict of interest.

---

**Figure S1a:**
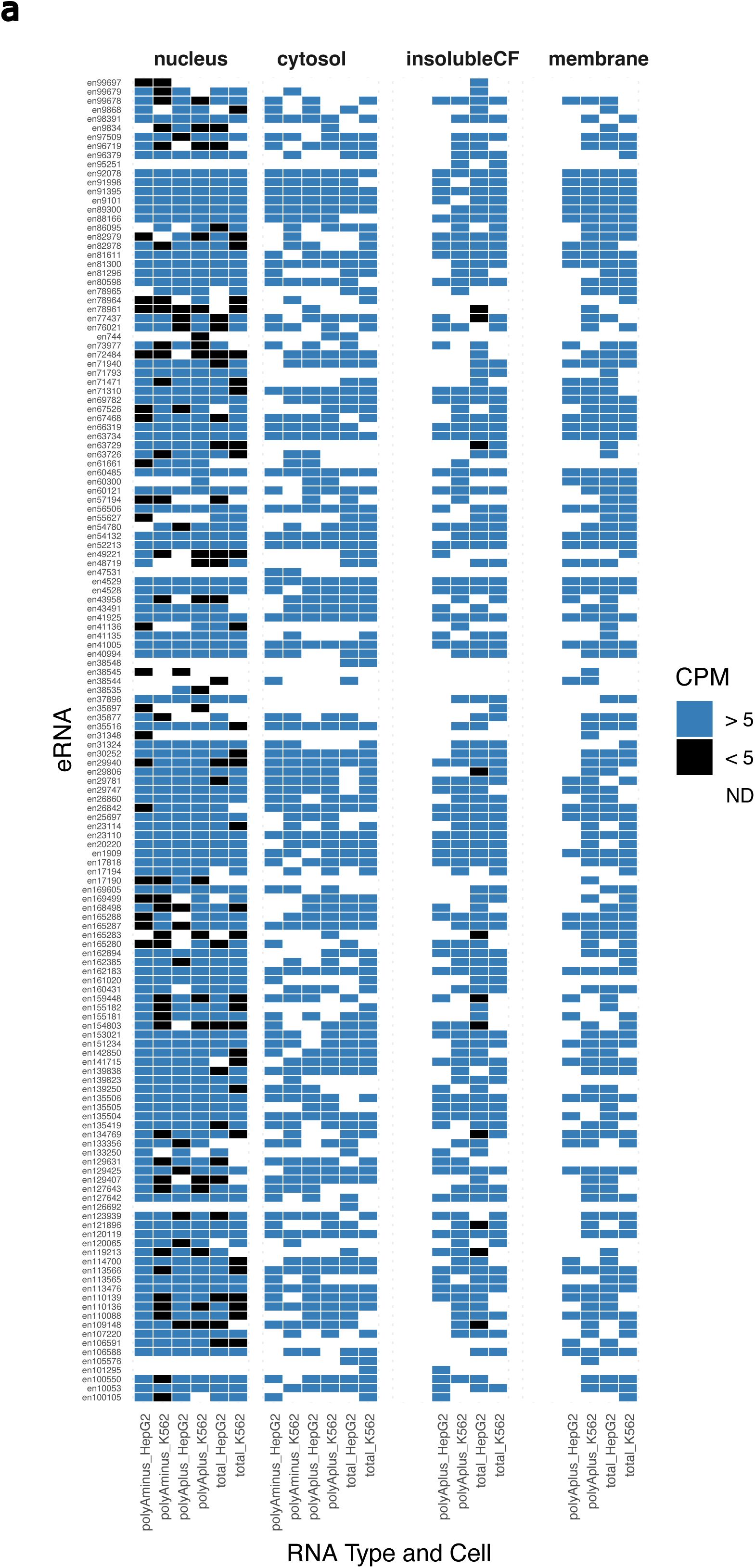
Heatmap of 186 regulatory eRNAs across subcellular fractions in K562 and HepG2 cells. eRNAs are grouped by subcellular location (nucleus, cytosol, insoluble cytoplasmic fraction, and membrane), and colour intensity reflects expression levels (CPM). Expression data were extracted from the eRNAkit resource.

**Figure S2a:**
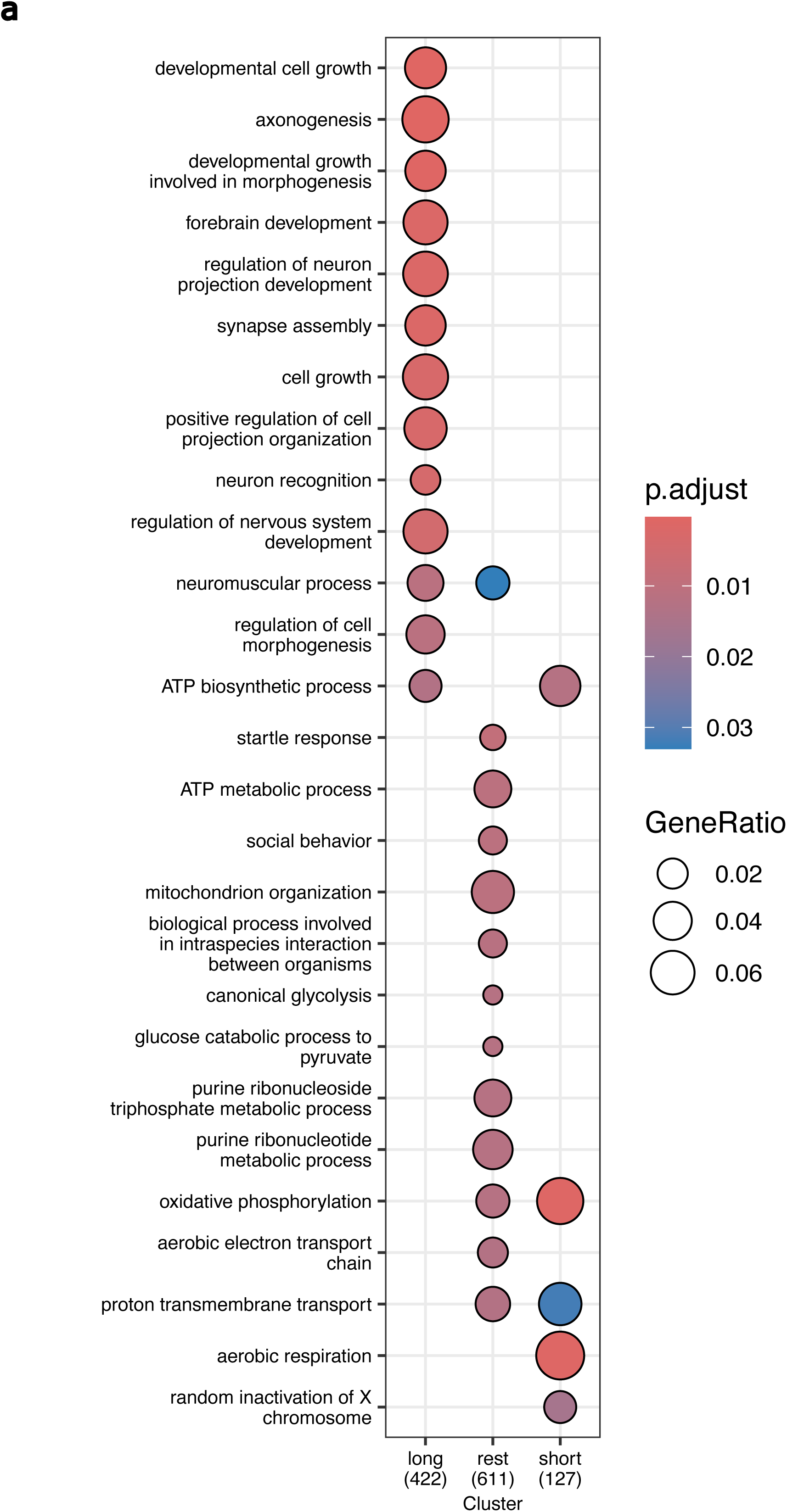
Gene ontology enrichment analysis of mRNAs targeted by eRNAs without known genomic interactions, grouped by distinct half-life classes (short, rest, and long). The top 12 enriched biological process terms are shown after reducing redundancy using semantic similarity filtering.

**Figure S3a-b:**
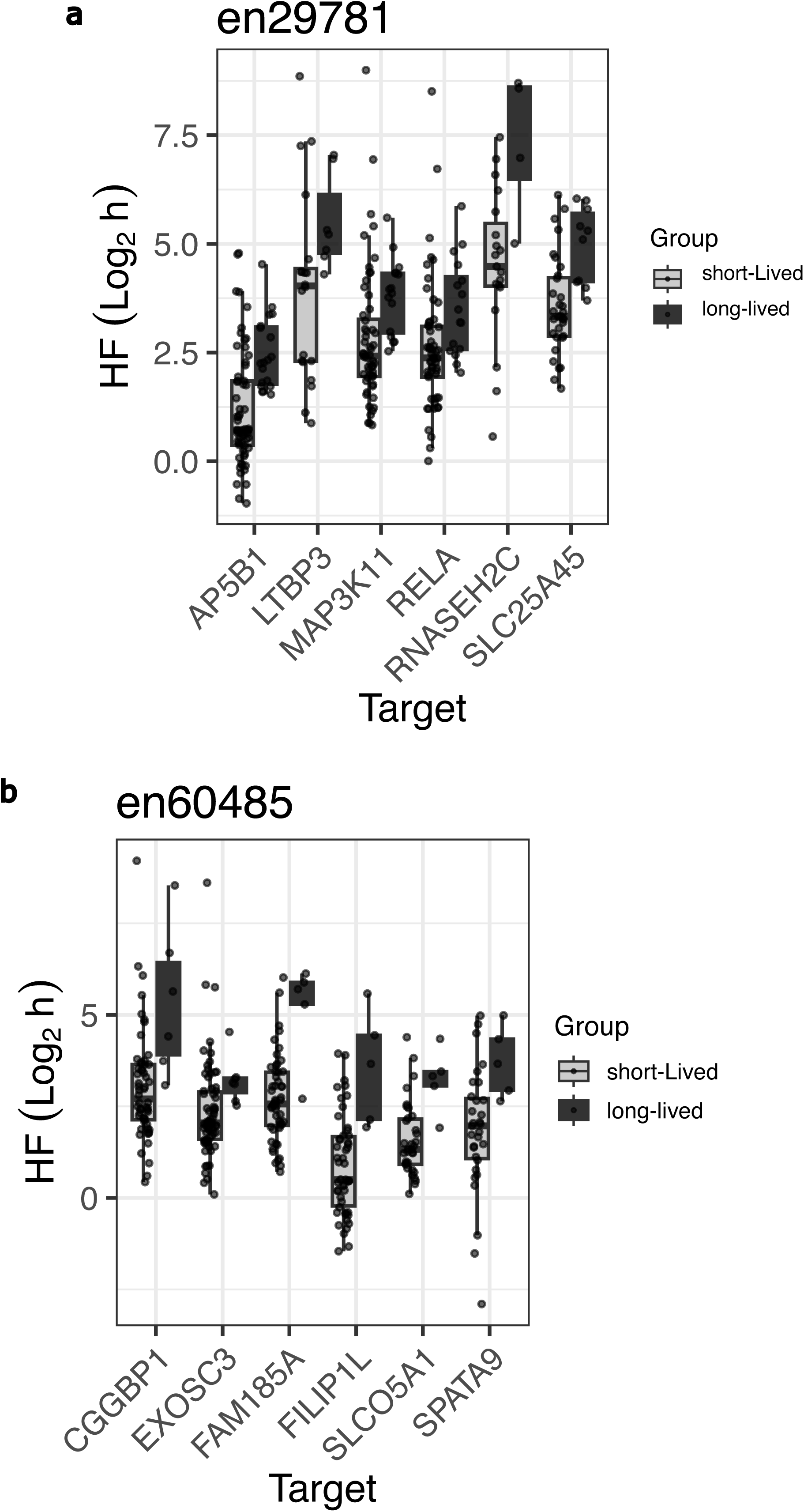
Comparison of mRNA half-lives for targets of (a) en29781 and (b) en60485 between sample groups classified by short-lived versus long-lived expression of en29781 or en60485 in the main time-course dataset, highlighting the most significant pairs.

**Figure S4a:**
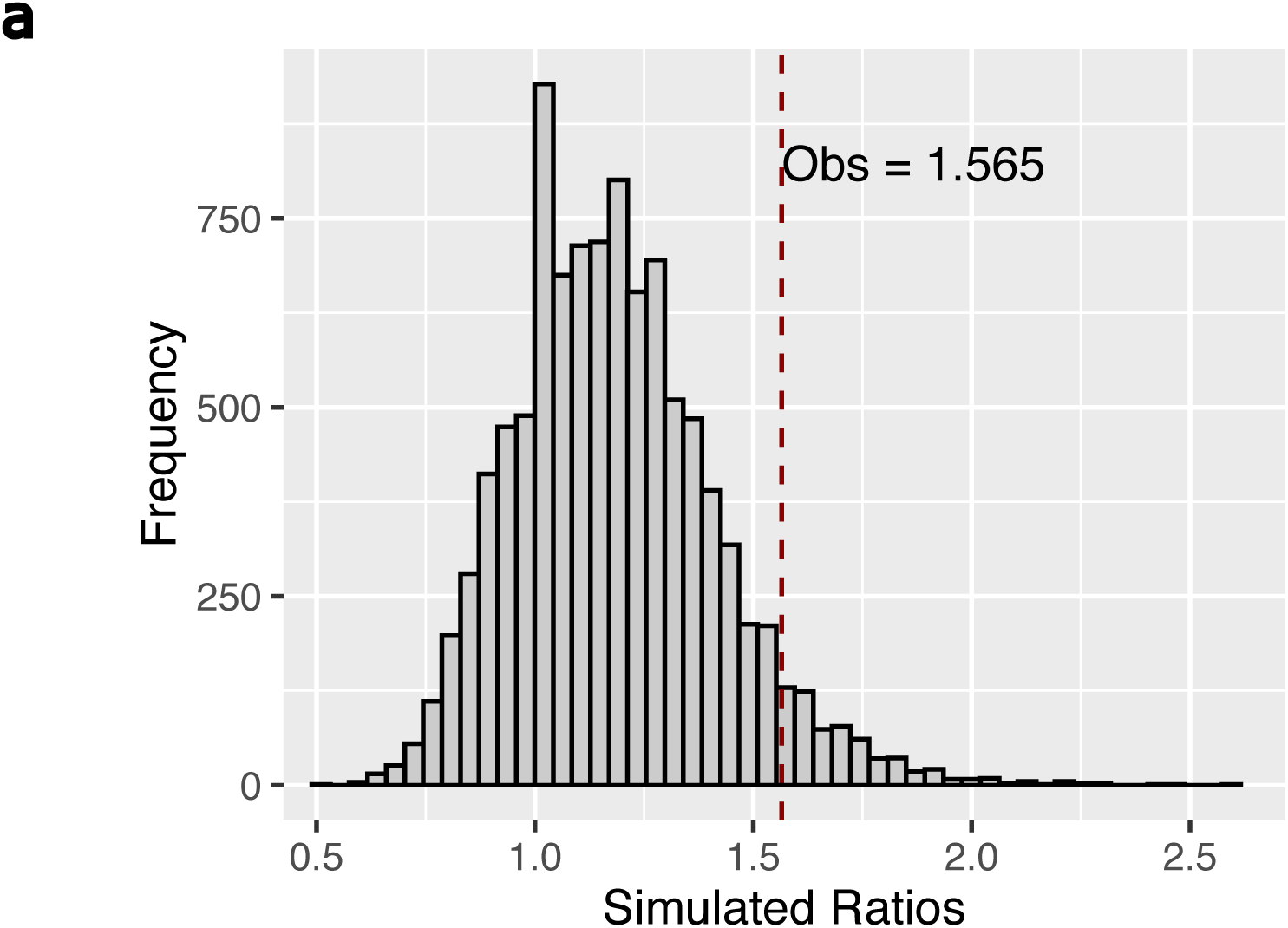
Distribution of overlap ratios from eRNA-mRNA pair shuffling simulations comparing regulatory eRNAs associated with half-life and translation efficiency (TE). The vertical line indicates the observed number of overlapping regulatory eRNAs.

## Notes

https://github.com/AneneLab/eRNAkit

### Competing Interest Statement

The authors have declared no competing interest.

## References

1. Kim, T.-K., Hemberg, M. & Gray, J. M. Enhancer RNAs: a class of long noncoding RNAs synthesized at enhancers. Cold Spring Harbor perspectives in biology 7, a018622 (2015).

2. Lewis, M. W., Li, S. & Franco, H. L. Transcriptional control by enhancers and enhancer RNAs. Transcription 10, 171–186 (2019).

3. Kim, T.-K. et al. Widespread transcription at neuronal activity-regulated enhancers. Nature 465, 182–187 (2010).

4. Arnold, P. R., Wells, A. D. & Li, X. C. Diversity and emerging roles of enhancer RNA in regulation of gene expression and cell fate. Frontiers in cell and developmental biology 7, 377 (2020).

5. Lam, M. T., Li, W., Rosenfeld, M. G. & Glass, C. K. Enhancer RNAs and regulated transcriptional programs. Trends in biochemical sciences 39, 170–182 (2014).

6. Benova, N., Kuklinkova, R., Eldahshoury, M. K. & Anene, C. A. eRNAkit: Expanding the Functional Atlas of human Enhancer RNAs Beyond the Nucleus. bioRxiv 2025–04 (2025).

7. Djebali, S. et al. Landscape of transcription in human cells. Nature 489, 101–108 (2012).

8. Sartorelli, V. & Lauberth, S. M. Enhancer RNAs are an important regulatory layer of the epigenome. Nature structural & molecular biology 27, 521–528 (2020).

9. Schwanhäusser, B. et al. Global quantification of mammalian gene expression control. Nature 473, 337–342 (2011).

10. Friedel, C. C., Dölken, L., Ruzsics, Z., Koszinowski, U. H. & Zimmer, R. Conserved principles of mammalian transcriptional regulation revealed by RNA half-life. Nucleic acids research 37, e115–e115 (2009).

11. Guhaniyogi, J. & Brewer, G. Regulation of mRNA stability in mammalian cells. Gene 265, 11–23 (2001).

12. Erdmann-Pham, D. D., Duc, K. D. & Song, Y. S. The key parameters that govern translation efficiency. Cell systems 10, 183–192 (2020).

13. Andersson, R. et al. An atlas of active enhancers across human cell types and tissues. Nature 507, 455–461 (2014).

14. Mikhaylichenko, O. et al. The degree of enhancer or promoter activity is reflected by the levels and directionality of eRNA transcription. Genes & development 32, 42–57 (2018).

15. Butcher, S. E. & Pyle, A. M. The molecular interactions that stabilize RNA tertiary structure: RNA motifs, patterns, and networks. Accounts of chemical research 44, 1302– 1311 (2011).

16. Kornienko, I. V., Aramova, O. Y., Tishchenko, A. A., Rudoy, D. V. & Chikindas, M. L. RNA Stability: A Review of the Role of Structural Features and Environmental Conditions. Molecules 29, 5978 (2024).

17. Wang, Y., Tang, Y., Xie, Z. & Wang, H. RPFdb v3. 0: an enhanced repository for ribosome profiling data and related content. Nucleic Acids Research 53, D293–D298 (2025).

18. Lennox, K. A. & Behlke, M. A. Cellular localization of long non-coding RNAs affects silencing by RNAi more than by antisense oligonucleotides. Nucleic acids research 44, 863–877 (2016).

19. Li, W., Notani, D. & Rosenfeld, M. G. Enhancers as non-coding RNA transcription units: recent insights and future perspectives. Nature Reviews Genetics 17, 207–223 (2016).

20. Rabani, M. et al. Metabolic labeling of RNA uncovers principles of RNA production and degradation dynamics in mammalian cells. Nature biotechnology 29, 436–442 (2011).

21. Tani, H. et al. Genome-wide determination of RNA stability reveals hundreds of short-lived noncoding transcripts in mammals. Genome research 22, 947–956 (2012).

22. Shi, H. et al. YTHDF3 facilitates translation and decay of N6-methyladenosine-modified RNA. Cell research 27, 315–328 (2017).

23. Huang, H. et al. Recognition of RNA N 6-methyladenosine by IGF2BP proteins enhances mRNA stability and translation. Nature cell biology 20, 285–295 (2018).

24. Collart, M. A. & Panasenko, O. O. The Ccr4–not complex. Gene 492, 42–53 (2012).

25. Hardy, E. C. & Balcerowicz, M. Untranslated yet indispensable—UTRs act as key regulators in the environmental control of gene expression. Journal of Experimental Botany 75, 4314–4331 (2024).

26. Slobodin, B. et al. Transcription dynamics regulate poly (A) tails and expression of the RNA degradation machinery to balance mRNA levels. Molecular cell 78, 434–444 (2020).

27. Zhang, Z. et al. An RNA tagging approach for system-wide RNA-binding proteome profiling and dynamics investigation upon transcription inhibition. Nucleic acids research 49, e65–e65 (2021).

28. Poetz, F. et al. RNF219 attenuates global mRNA decay through inhibition of CCR4-NOT complex-mediated deadenylation. Nature Communications 12, 7175 (2021).

29. Ma, X. et al. N 6-methyladenosine modification-mediated mRNA metabolism is essential for human pancreatic lineage specification and islet organogenesis. Nature Communications 13, 4148 (2022).

30. Block, C. J. et al. RNA binding protein RBMS3 is a common EMT effector that modulates triple-negative breast cancer progression via stabilizing PRRX1 mRNA. Oncogene 40, 6430–6442 (2021).

31. Sharma, S. et al. Acetylation-dependent control of global Poly (A) RNA degradation by CBP/p300 and HDAC1/2. Molecular cell 63, 927–938 (2016).

32. Lim, J. et al. Uridylation by TUT4 and TUT7 marks mRNA for degradation. Cell 159, 1365–1376 (2014).

33. Poetz, F. et al. Control of immediate early gene expression by CPEB4-repressor complex-mediated mRNA degradation. Genome Biology 23, 193 (2022).

34. Slobodin, B. et al. Cap-independent translation and a precisely located RNA sequence enable SARS-CoV-2 to control host translation and escape anti-viral response. Nucleic acids research 50, 8080–8092 (2022).

35. Zhao, Z. et al. QKI shuttles internal m7G-modified transcripts into stress granules and modulates mRNA metabolism. Cell 186, 3208–3226 (2023).

36. Xu, Z. et al. KHSRP stabilizes m6A-modified transcripts to activate FAK signaling and promote pancreatic ductal adenocarcinoma progression. Cancer Research 84, 3602– 3616 (2024).

37. Sung, H.-M. et al. Stress-induced nuclear speckle reorganization is linked to activation of immediate early gene splicing. Journal of Cell Biology 222, e202111151 (2023).

38. Quarto, G. et al. Fine-tuning of gene expression through the Mettl3-Mettl14-Dnmt1 axis controls ESC differentiation. Cell (2025).

39. Chen, C.-Y. A., Zhang, Y., Xiang, Y., Han, L. & Shyu, A.-B. Antagonistic actions of two human Pan3 isoforms on global mRNA turnover. Rna 23, 1404–1418 (2017).

40. Lugowski, A., Nicholson, B. & Rissland, O. S. DRUID: a pipeline for transcriptome-wide measurements of mRNA stability. Rna 24, 623–632 (2018).

41. Vicente, C. et al. The CCR4-NOT complex is a tumor suppressor in Drosophila melanogaster eye cancer models. Journal of Hematology & Oncology 11, 1–16 (2018).

42. Wang, X. et al. N 6-methyladenosine-dependent regulation of messenger RNA stability. Nature 505, 117–120 (2014).

43. Murakawa, Y. et al. RC3H1 post-transcriptionally regulates A20 mRNA and modulates the activity of the IKK/NF-κB pathway. Nature communications 6, 7367 (2015).

44. Choe, J. et al. mRNA circularization by METTL3–eIF3h enhances translation and promotes oncogenesis. Nature 561, 556–560 (2018).

45. Gao, X. et al. Low RNA stability signifies strong expression regulatability of tumor suppressors. Nucleic acids research 51, 11534–11548 (2023).

46. Huang, H. et al. Histone H3 trimethylation at lysine 36 guides m6A RNA modification co-transcriptionally. Nature 567, 414–419 (2019).

47. Kapeli, K. et al. Distinct and shared functions of ALS-associated proteins TDP-43, FUS and TAF15 revealed by multisystem analyses. Nature communications 7, 12143 (2016).

48. Liu, C. et al. IGF2BP3 promotes mRNA degradation through internal m7G modification. Nature Communications 15, 7421 (2024).

49. Cogan, J. A., Benova, N., Kuklinkova, R., Boyne, J. R. & Anene, C. A. Meta-analysis of RNA interaction profiles of RNA-binding protein using the RBPInper tool. Bioinformatics Advances 4, vbae127 (2024).

50. Bolger, A. M., Lohse, M. & Usadel, B. Trimmomatic: a flexible trimmer for Illumina sequence data. Bioinformatics 30, 2114–2120 (2014).

51. Kim, D., Langmead, B. & Salzberg, S. L. HISAT: a fast spliced aligner with low memory requirements. Nature methods 12, 357 (2015).

52. Anders, S., Pyl, P. T. & Huber, W. HTSeq—a Python framework to work with high-throughput sequencing data. Bioinformatics 31, 166–169 (2015).

53. Yu, G., Wang, L.-G., Han, Y. & He, Q.-Y. clusterProfiler: an R package for comparing biological themes among gene clusters. Omics: a journal of integrative biology 16, 284–287 (2012).

54. Wang, J. Z., Du, Z., Payattakool, R., Yu, P. S. & Chen, C.-F. A new method to measure the semantic similarity of GO terms. Bioinformatics 23, 1274–1281 (2007).

55. Anene, C., Graham, A. M., Boyne, J. & Roberts, W. Platelet microparticle delivered microRNA-Let-7a promotes the angiogenic switch. Biochimica et Biophysica Acta (BBA)-Molecular Basis of Disease 1864, 2633–2643 (2018).

